# Immunophenotyping and transcriptional profiling of human plasmablasts in dengue

**DOI:** 10.1101/2021.04.09.439257

**Authors:** Charu Aggarwal, Keshav Saini, Elluri Seetharami Reddy, Mohit Singla, Kaustuv Nayak, Yadya M. Chawla, Deepti Maheshwari, Prabhat Singh, Pragati Sharma, Priya Bhatnagar, Sanjeev Kumar, Kamalvishnu Gottimukkala, Harekrushna Panda, Sivaram Gunisetty, Carl W Davis, Haydn Thomas Kissick, Sushil Kumar Kabra, Rakesh Lodha, Guruprasad R Medigeshi, Rafi Ahmed, Kaja MuraliKrishna, Anmol Chandele

## Abstract

Previous studies have shown that plasmablasts expand massively in dengue patients as compared to many other situations such as influenza infection or vaccination. However, a detailed understanding of the phenotypes and transcriptional features of these cells is lacking. Moreover, despite India having nearly a third of global dengue disease burden, there is virtually no information on plasmablasts responses in dengue patients from India. Here, we provide a detailed characterization of plasmablast responses from dengue confirmed febrile children in India. Immunophenotyping and RNA seq analysis showed that in addition to secreting dengue specific antibodies, these massively expanding cells expressed several adhesion molecules, chemokines and chemokine receptors that are involved in endothelial interactions, homing to skin or mucosal tissues including intestine. Surprisingly, we found that these cells also upregulated expression of several cytokine genes that are involved in angiogenesis, leukocyte extravasation and vascular permeability. These transcriptional features were qualitatively similar to plasmablasts from influenza vaccinees. Interestingly, the expansion of the plasmablasts in dengue patients was significantly lower in patients with primary dengue infection compared to those with secondary dengue. Moreover, within the primary dengue patients, their expansion was significantly lower in patients with mild dengue infection (DI) compared to patients with dengue with warning signs (DW) or severe dengue (SD). These results significantly improve our understanding of human plasmablast responses in dengue.

Importance

Dengue is a globally spreading with over 100 million clinical cases annually with symptoms ranging from mild self-limiting febrile illness to more severe and sometimes life-threatening dengue hemorrhagic fever or shock, especially among children. India contributes nearly a third of global dengue disease burden. The pathophysiology of dengue is complex and remains poorly understood despite many advances indicating a key role for antibody dependent enhancement of infection. While serum antibodies have been extensively studied, the characteristics of the cellular factories responsible for antibody production, i.e., plasmablasts, are only beginning to emerge. This study provides a comprehensive understanding of the magnitude, phenotype, functional and transcriptional profiles of human plasmablasts from dengue patients in India.

## Introduction

Dengue is emerging as one of the most dangerous mosquito-borne human hemorrhagic fever virus disease. Worldwide, over 100 million clinically significant cases are estimated to occur each year (1). This acute systemic infection results in clinical disease ranging from mild self-limiting febrile illness to more severe and sometimes life-threatening dengue hemorrhagic fever (DHF) or dengue shock syndrome (DSS), especially among children (2, 3). While the precise mechanisms of this varied disease spectrum are still being elucidated, numerous studies suggest immune components are involved in these pathological manifestations. This is evidenced by studies showing an association of DHF/DSS with a massive T and B cell response (4–6); an accumulation of several inflammatory cytokines implicated in vascular leakage in patients manifesting DHF/ DSS (7–10); and an increased disease susceptibility in individuals who are previously exposed to a heterologous dengue serotype (11–13). Additionally, while neutralizing antibodies protect; sub-neutralizing or non-neutralizing antibodies are believed to contribute to an increased viral uptake and replication in cells via Fc receptors through a process called antibody dependent enhancement of infection (ADE) (14, 15).

While antibody responses to dengue have been extensively studied (16–21), a detailed understanding of the cells responsible for producing these antibodies during the acute febrile phase, i.e., the plasmablasts or antibody secreting cells (ASCs), is only beginning to emerge. These emerging studies have revealed that plasmablasts expand rapidly and massively and usually peak around the time of subsidence of fever and serious symptoms (5, 22–24). However, there is very limited information on the transcriptional features of the human plasmablasts beyond antibody production in dengue (25–29). Moreover, despite India contributing to nearly a third of the global dengue disease burden, thus far, virtually no information is available about plasmablasts in dengue patients from India,. Considering these factors, in this study we characterized expansion, functional phenotypes, and transcriptional profiles of the plasmablasts from dengue confirmed febrile children in India.

## Results

### Characteristics of the dengue patients included in the study

This study is based on dengue confirmed febrile children recruited from All India Institute of Medical Sciences (AIIMS), New Delhi through years 2012-2019. A total of 199 children in whom sufficient blood sample was obtained for plasmablast enumeration are included in this analysis. The patient cohort was comprised of both male and female children with ages ranging between 1.2 to 15 years **(Table 1**). Average day of symptoms was 4.5. A blood sample was collected at the time of patient recruitment and where possible a month later (convalescent). At the time of recruitment, 45 patients had mild dengue infection without warning signs (DI),75 patients had dengue with warning signs (DW), and 79 patients had severe dengue (SD). The patients were comprised of a substantial mix of primary and secondary dengue infections.

**Table 1.** Characteristics of the dengue patient cohort.

|  |  |
| --- | --- |
| Male/Female (n) | 105/94 |
| Age in years, average (Range) | 9.3 (1.2-15) |
| Day of symptoms, average (Range) | 4.5 (2–7) |
| Severity* |  |
| Dengue Infection (DI) | 45 |
| Dengue Warning (DW) | 75 |
| Severe Dengue (SD) | 79 |
| Serological status** |  |
| Primary | 81 |
| Secondary | 118 |

\*Disease grade classification was done by WHO 2009 criteria (96)
\*\* Serological status determined by WHO classification using Panbio Dengue IgM and IgG Elisa (96)

### Immunophenotyping of the plasmablasts from dengue patients

Although many studies from other parts of the world showed plasmablasts expand massively in dengue patients (5, 22–24), given that there is virtually no information on plasmablast responses in dengue patients from India. Thus, we first examined plasmablast expansion in our patient cohort. Gating strategy for enumeration of plasmablasts in peripheral blood derived mononuclear cells (PBMC) is shown in **Fig. 1A**. Plasmablasts were identified within the gated CD19^+^ B cells that were strongly positive for CD27 and CD38, with a low or no CD20 expression. Consistent with previous studies from other parts of the world (5, 22–24), we found that plasmablasts showed a massive and heterogenous expansion in these patients with frequencies reaching as high as 80% of the total B cell population in some patients (**Fig. 1B)**. These responses appeared transiently during the acute febrile phase of dengue. We then asked what are the phenotypic and functional features of these massively expanding plasmablasts. A detailed immunophenotyping of the plasmablasts is shown in **Fig. 1C**. Consistent with their definition, we found that the plasmablasts were blasting as evidenced by high forward scatter (FSC) and side scatter (SSC). They were highly proliferating as indicated by expression of proliferating cell nuclear antigen Ki67, transferrin receptor CD71; and downregulation of heat stable antigen CD24. They expressed little or no CD20, a key molecule involved in efficient signaling and calcium mobilization following ligation of the B cell receptor (BCR) or other surface proteins of B cells (30, 31).

**FIG 1.**
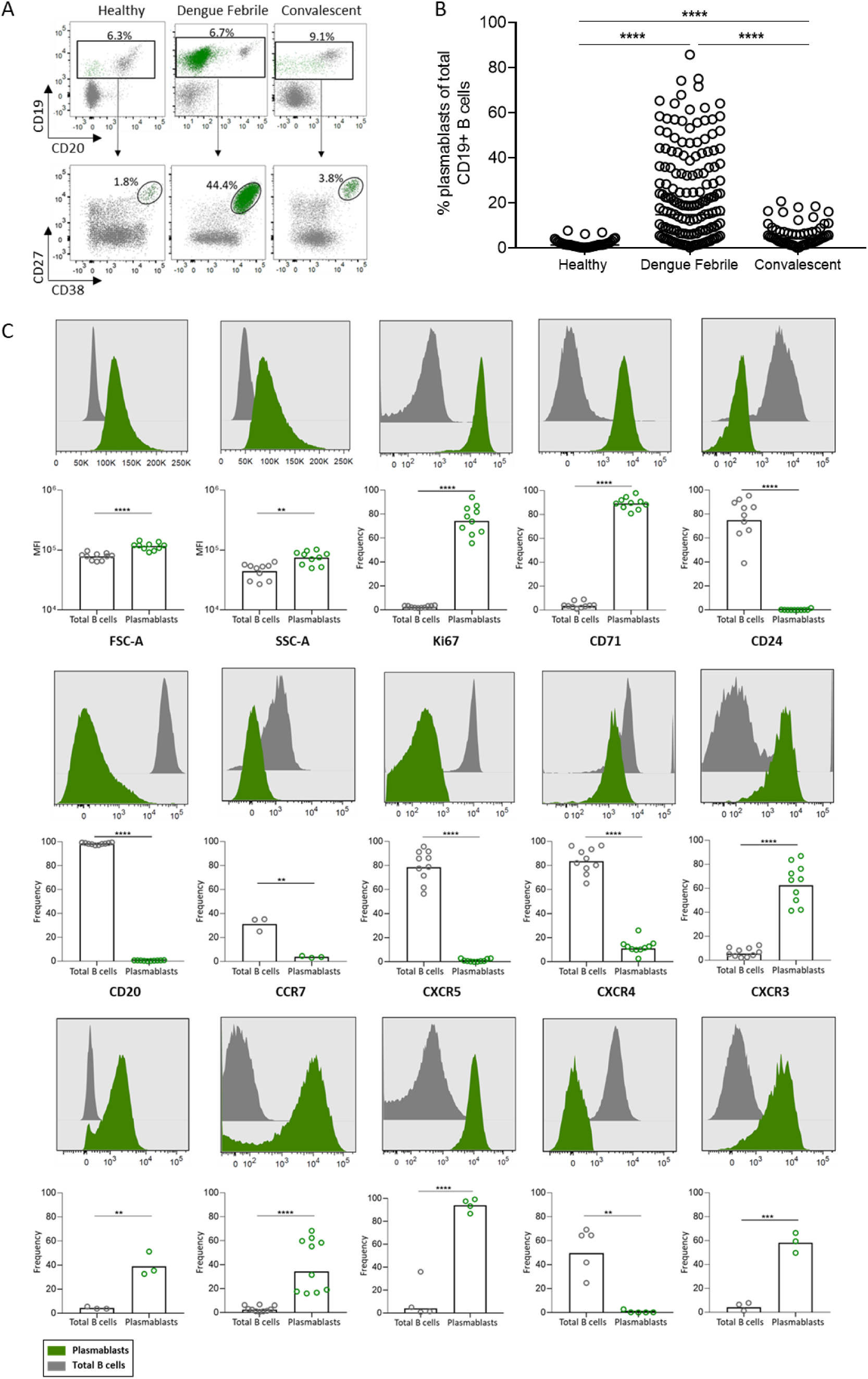
Immunophenotyping of the plasmablasts from dengue patients. (A) Plots showing gating strategy for identification of plasmablasts. Top panel shows the B cells among the total PBMCs (CD19+CD20+ cells) and bottom panel shows the plasmablasts (CD27+CD38+ cells) identified among the gated B cell population in healthy subject, acute dengue febrile patient and dengue recovered patient (convalescent). The green colored population in each plot represents the plasmablasts. (B) Scatter dot plot showing plasmablast frequencies of total CD19+ B cells in healthy subject (n=43), acute dengue febrile patient (n=162) and convalescent subject (n=68). The line represents the geometric mean of all points. (C) Phenotype of plasmablasts (green) and other B cells (grey) is shown as overlay histograms. Corresponding box and scatter plots for each of the parameters analyzed is shown below the overlay histogram. Each dot represents a single individual.

Further analysis of chemokine receptor expression showed that the plasmablasts robustly downregulated germinal center homing chemokine receptors CCR7 and CXCR5 as well as the bone-marrow homing chemokine receptor CXCR4 - suggesting that these cells are destined to circulate in the periphery rather than migrate to primary or secondary lymphoid tissues (bone marrow/ germinal centers). By contrast they robustly upregulated chemokine receptors CXCR3 and CCR2, implying that they are equipped with ability to home to inflamed tissues. Interestingly, they also upregulated cutaneous lymphocyte associated antigen (CLA) indicating they acquire the ability to interact with vascular endothelial cells via endothelial leukocyte adhesion molecule-1 (ELAM1) and infiltrate cutaneous tissue. They also robustly expressed CCR9, suggesting that they are equipped with ability to home to not only cutaneous tissue but also to other mucosal/ intestinal tissues.

Analysis of key cytokine receptors revealed that plasmablasts downregulated B cell activating factor receptor, BAFF-R, while upregulating receptors for interleukin-6 (IL6R). This suggested that these cells are likely to be inept in receiving pro-survival signals through the BAFF-BAFF-R axis but have acquired the ability to efficiently receive signals from other pro-survival cytokines such as IL-6 that are systemically available during the acute febrile phase of dengue infection (9, 32, 33).

Consistent with the appreciation that plasmablasts are primarily antibody secreting cells, we also found that they expressed little or no surface IgD, IgM or IgG, but were robustly positive for intracellular expression of IgG or IgM or IgA **(Fig. 2A)**. Analysis of antibody secreting cell (ASC) functions by functional *ex vivo* ELISpot assays revealed that these plasmablasts were robustly secreting IgG or IgM and the total ASC response remarkably reflected dengue specific ASC response during the acute febrile phase **(Fig. 2B)**.

**FIG 2.**
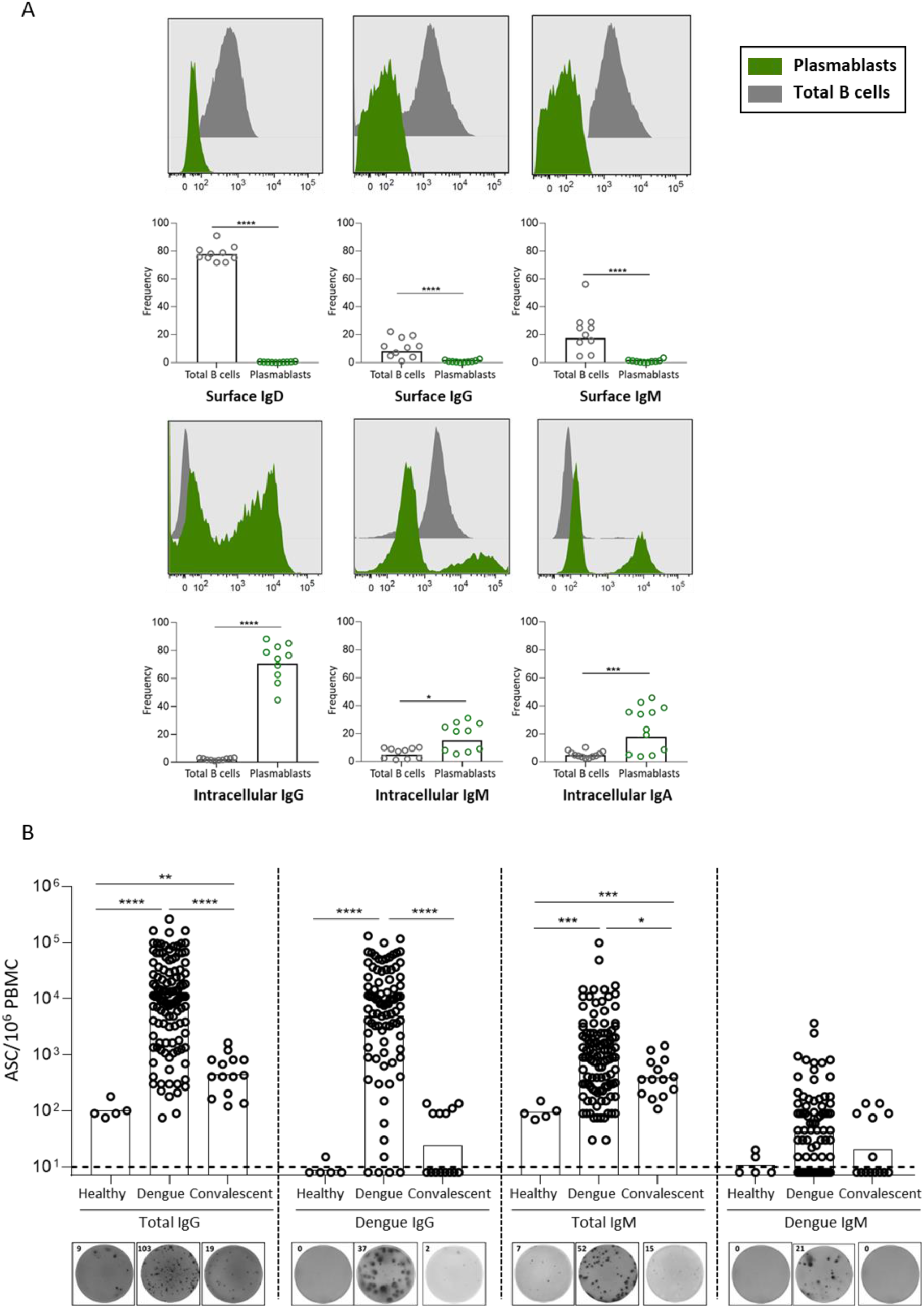
Antibody secreting function of the plasmablasts from dengue patients. (A) Top panel, Histograms (top) and scatter plots showing (bottom) surface antibody markers (IgD, IgG and IgM) in plasmablasts (green) and total B cells (grey) in dengue patients. Bottom panel, Histograms (top) and scatter plots showing (bottom) showing intracellular IgG, IgM and IgA in plasmablasts (green) and total B cells (grey) in dengue patients. (B) Box and scatter plot (top) showing the frequencies total IgG ASC/million PBMC (leftmost), dengue specific IgG ASC/million PBMC (second from left), total IgM ASC/million PBMC (third from the left) and dengue specific IgM ASC/million PBMC (rightmost) in healthy subject (n=6), acute dengue febrile patient (n=118) and convalescent subject (n=14). Corresponding ELISpot pictures (bottom) showing the total and dengue specific IgG and IgM spots. The numbers on the top left corner of the ELISpot pictures represent the number of spots counted in a well.

Taken together, these results show that these massively proliferating plasmablasts that transiently appear in blood circulation during dengue febrile phase are conditioned to receive pro-survival signals through cytokines that are systemically available during the acute febrile phase of dengue. Moreover, in addition to secreting dengue-specific antibodies, these cells are also equipped with ability to interact with endothelial cells and home to inflamed tissues, mucosa, intestine and/or skin.

### Transcriptional profiling of the plasmablasts from dengue patients

Although many studies have conducted transcriptional profiling of plasma cells (25, 26, 29, 34- 39), transcriptional profiling of human plasmablasts is surprisingly scarce (25–29). Also, so far there is no information on the global gene expression profiles of plasmablasts from dengue patients although one study that performed single cell RNA seq of the plasmablasts from dengue patients only analyzed immunoglobulin gene usage (27). To gain a detailed understanding of the plasmablasts beyond antibody production and the phenotypic analysis described above, we sorted plasmablasts from dengue patients and performed a global transcriptomic analysis using bulk RNA sequencing (GSE171487). The characteristics of the patients from whom the plasmablasts were sorted are listed in **Table 2**. Naïve B cells sorted simultaneously from the same patients were included for comparison. The sort strategy is shown in **Fig. 3**.

**FIG 3.**
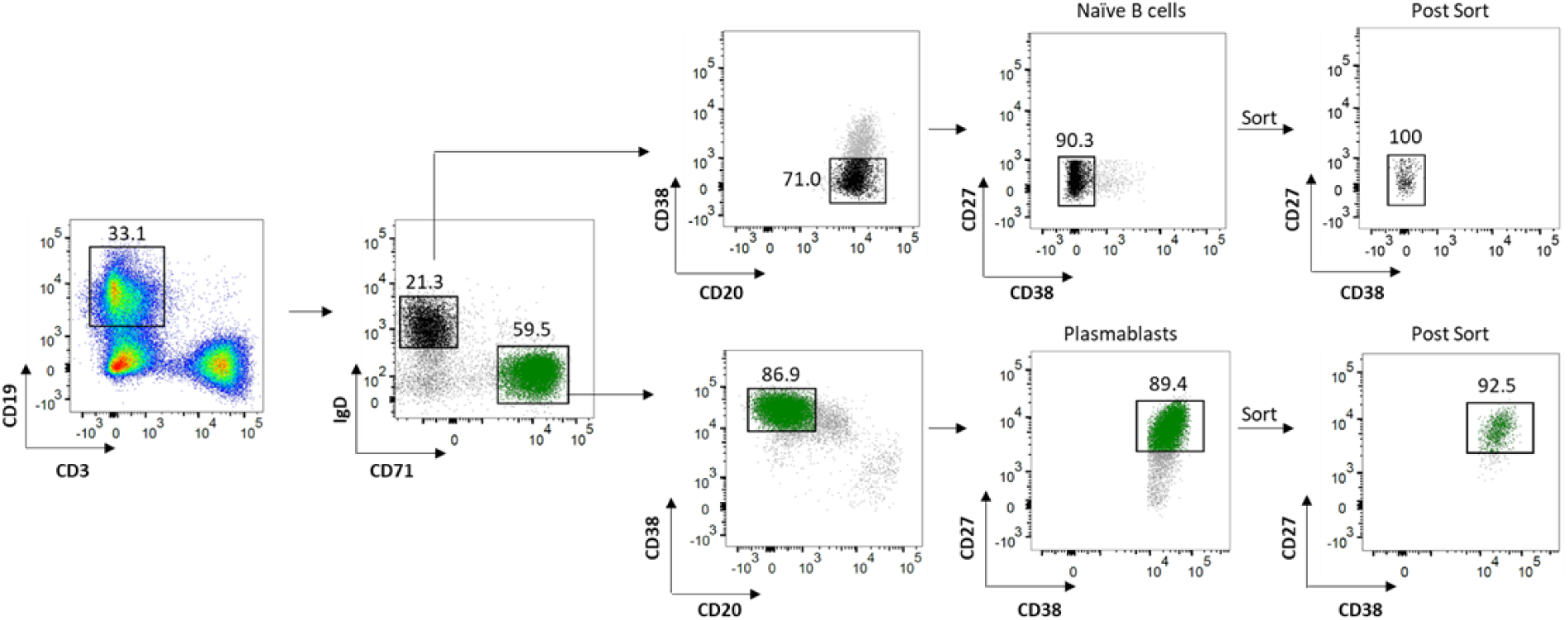
Sorting strategy for the naïve B cells and plasmablasts. The PBMCs were stained as described in the methods. Live blasting CD19+ B cells were gated to get IgD+CD71- (black) and IgD-CD71+ B cells (green). They were further gated to get CD20+ CD27- CD38- naïve (black, top panel) B cells and CD20- CD27+ CD38+ plasmablasts (green, bottom panel) respectively and sorted with high purity. The numbers represent the frequencies of gated population with the rightmost plot depicting the frequencies of naïve B cells and plasmablasts after sorting.

**Table 2.** Characteristics of dengue patients from whom plasmablasts and naïve B cells were sorted.

| Subject# | Male (M)/Female (F) | Primary (P)/Secondary (S)<br>status | Disease Severity |
| --- | --- | --- | --- |
| 1 | F | P | SD |
| 2 | M | P | SD |
| 3 | F | P | SD |
| 4 | F | S | SD |
| 5 | M | P | DW |
| 6 | F | S | DW |
| 7 | F | S | DW |
| 8 | M | S | DW |

Because plasmablasts are primarily antibody secreting cells and thus the bulk of their RNA is expected to be of immunoglobulin genes, we first asked what portion of the transcripts accounted for immunoglobulin (Ig) genes. Of the 98.4 million reads of plasmablasts, 27.9% of total reads (27.5 million) was represented by 407 Ig genes, confirming that Ig gene transcription was highly active in plasmablasts as compared to naïve B cells. By contrast, the Ig gene reads contributed only 2.8% (3.7 million) of total reads of naïve B cells (129.6 million). A total of 130 Ig genes that were differentially expressed in plasmablasts compared to naïve B cells are shown in **Fig.4**. Heavy chains of the IgD, IgE, IgM, IgA and IgG subtypes are marked by boxes; and consistent with our immunophenotyping analysis, plasmablasts substantially down-regulated IgD; and upregulated IgG, IgA and IgM.

**FIG 4.**
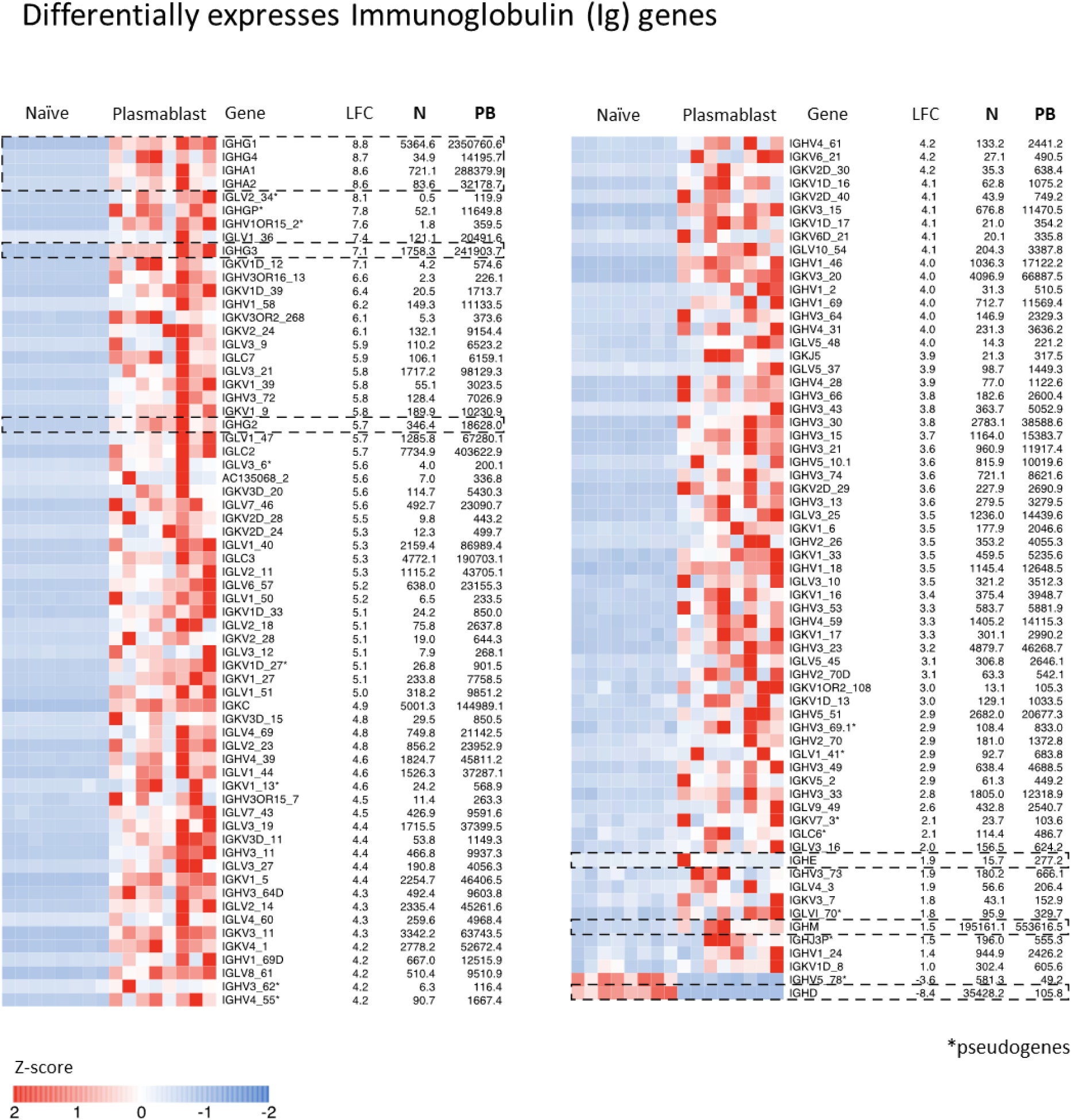
Heat maps of the differentially expressed Immunoglobulin genes. 130 immunoglobulin genes including variable and constant regions of heavy and light chains and pseudogenes are shown in the heatmap. IgG, IgA, IgM, IgE and IgD are shown in boxes. Pseudogenes are marked by an asterisk (*). Gradient of high to low gene expression based on z-score normalized counts is indicated from red to blue color. Genes that were significantly differential were only considered and clustered subjects using ward.D2 method. Mentioned on right side of heatmap are gene name with common name (in brackets), log2 fold change (LFC), average normalized counts of naïve (N) and average normalized counts of plasmablast (PB).

Principal component analysis (PCA) of plasmablasts versus naïve B cells showed that plasmablasts had a highly distinct transcriptional profile compared to naïve B cells even after excluding the Ig reads **(Fig. 5A)**. A total of 4137 non-Ig genes were differentially expressed in plasmablasts compared to naïve B cells, as shown in volcano plot (**Fig. 5B**). Of these, 2062 genes were upregulated and 2075 genes were downregulated in plasmablasts compared to naïve B cells. A heatmap of these differentially expressed genes is shown in **Fig. 5C**, and the entire set is provided in **Supplementary Table 1** (GSE171487). Interestingly, although the plasmablasts clustered separately from naïve B cells, there was no distinct clustering observed for plasmablasts from primary / secondary or from dengue patients with warning signs (DW) or with severe disease (SD). Analysis of select genes that are typically expected to be upregulated in plasmablasts (*XBP1, CD27, PRDM1, CD38, CXCR3 and IRF4*) and select genes that are typically expected to be downregulated in plasmablasts (*CD40, BCL6, ID3, CXCR4, EBF1, PAX5, IRF8, SPIB, BACH2, CD22, MS4A1, CD24 and CXCR5*) showed expected pattern thereby validating our data (**Fig. 5D**). Additionally, we found that our immuno-phenotyping profiles presented in the section above closely mirrored our RNA Seq transcriptional profiles (**Fig. 5E).**

**FIG 5.**
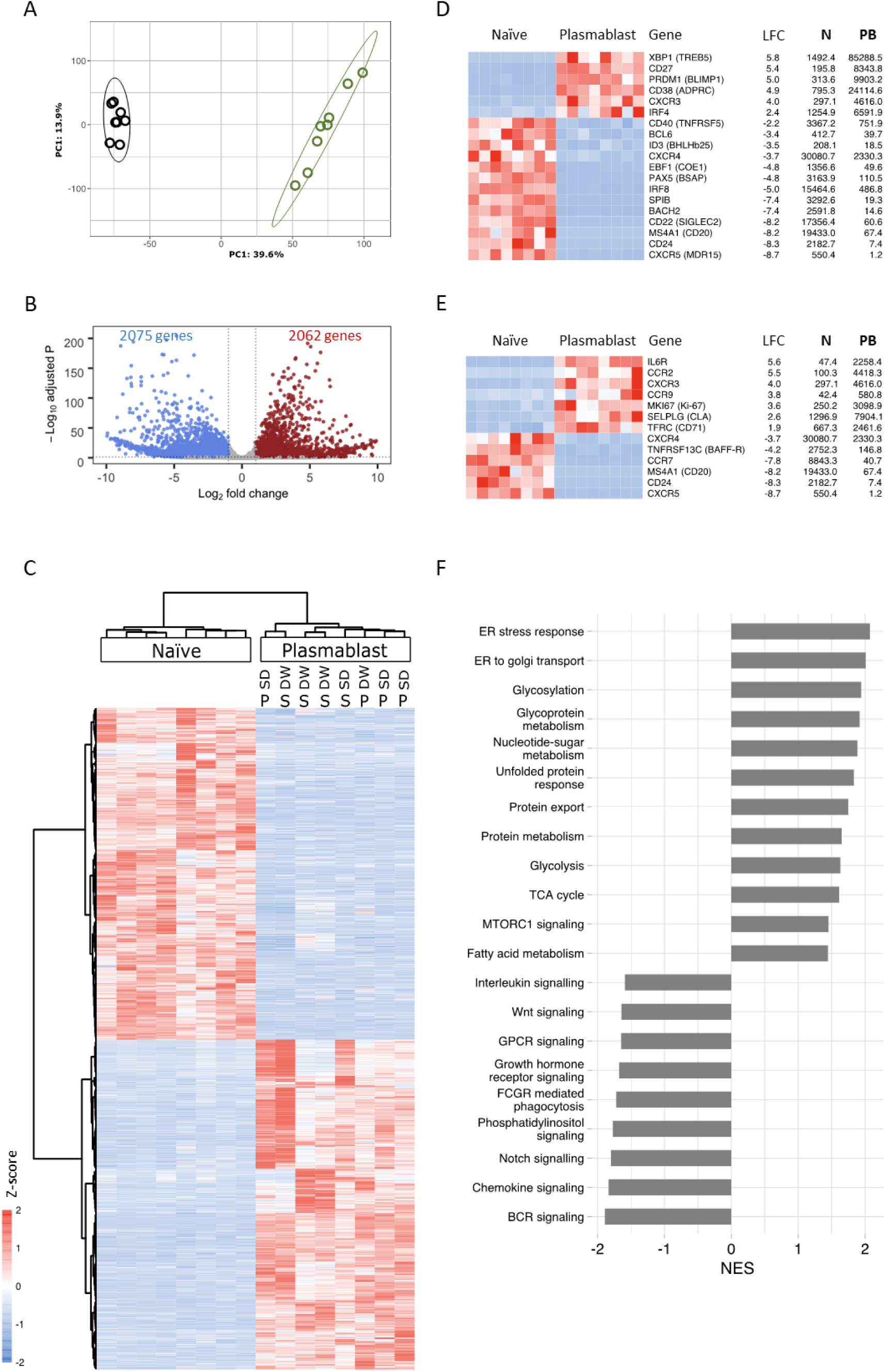
Transcriptional profiling of the plasmablasts from dengue patients. (A) Principal component analysis (PCA) score plot of normalized counts of 14,773 genes showing clustering within plasmablasts (green) and naïve B cells (black) from eight different acute dengue febrile subjects. (B) Volcano plot with -log (adjusted P value) on y-axis and log2 (fold change) on x-axis. Scattered dots represent genes. Red dots are genes that are significantly upregulated and blue dots represent significantly downregulated genes in plasmablasts versus naïve B cells (total: 4137 genes), whereas grey dots are either non-significant, non-differential or both. (for *significant*: adjusted P value < 0.05 and P value < 0.01; for *differential*: log2 fold change >= 1 or =<-1). (C) Heatmap showing the 4137 significantly upregulated (red) or downregulated (blue) genes in plasmablasts compared to naïve B cells from eight different acute dengue febrile patients. Z-score of normalized counts were taken for the heatmap. Ward.D2 method was used for clustering. (for *significant*: adjusted P value < 0.05 and P value < 0.01; for *differential*: log2 fold change >= 1 or =<-1). (D) Gene expression profile of select genes that are already known in literature, and (E) immunophenotypic markers used in flow cytometric analysis for plasmablasts. Red- highest and blue- lowest. The gene name with common name in brackets, log2 fold change (LFC), average normalized counts of naïve (N) and average normalized counts of plasmablast (PB) are indicated below the heatmap and are arranged as per the descending LFC values. Z-score of normalized counts were taken for the heatmap. (F) Gene set enrichment analysis (GSEA) performed using gene sets derived from GO: biological process, Hallmark, KEGG and Reactome gene sets from MSigDB. Significant pathways with FDR q value < 25% are shown. Enriched term names are manually collapsed. Positive and negative NES correlates to upregulated and downregulated pathways, respectively in plasmablasts compared to naïve B cells.

Gene set enrichment analysis (GSEA) of the differentially expressed genes is shown in **Supplementary Table 2**. Of the top 20 enriched terms for upregulated genes in plasmablasts, processes related to cellular metabolism, protein synthesis, glycosylation, transport and secretion were highly significant (FDR <25%) **(Fig. 5F).** By contrast, the down regulated genes were associated with processes related to signaling through B cell receptor (BCR) or signaling through other external ligands such as notch, phosphatidyl inositol, growth hormone receptor, G protein coupled receptors, WNT and interleukins. This pattern is consistent with the idea that these plasmablasts are highly differentiated short-lived antibody producing effector cells that become largely refractory to BCR signaling or other external stimuli. The differentially expressed genes in each of the biological processes is provided as an extensive list in **Supplementary Table 3**. Additionally, heatmaps of the differentially expressed genes associated with proliferation, metabolism, antibody synthesis, glycosylation, growth factors, transcription factors and Interferon stimulating genes (ISGs), their log2fold change (LFC), and average counts of naïve (N) and plasmablast (PB) are shown in **Fig.6**. Taken together, these data suggest that plasmablasts are highly proliferating, metabolically active, cellular factories specialized with antibody synthesis, glycosylation and secretion, while downregulating signaling through B cell receptors or through many other external stimuli.

**FIG 6.**
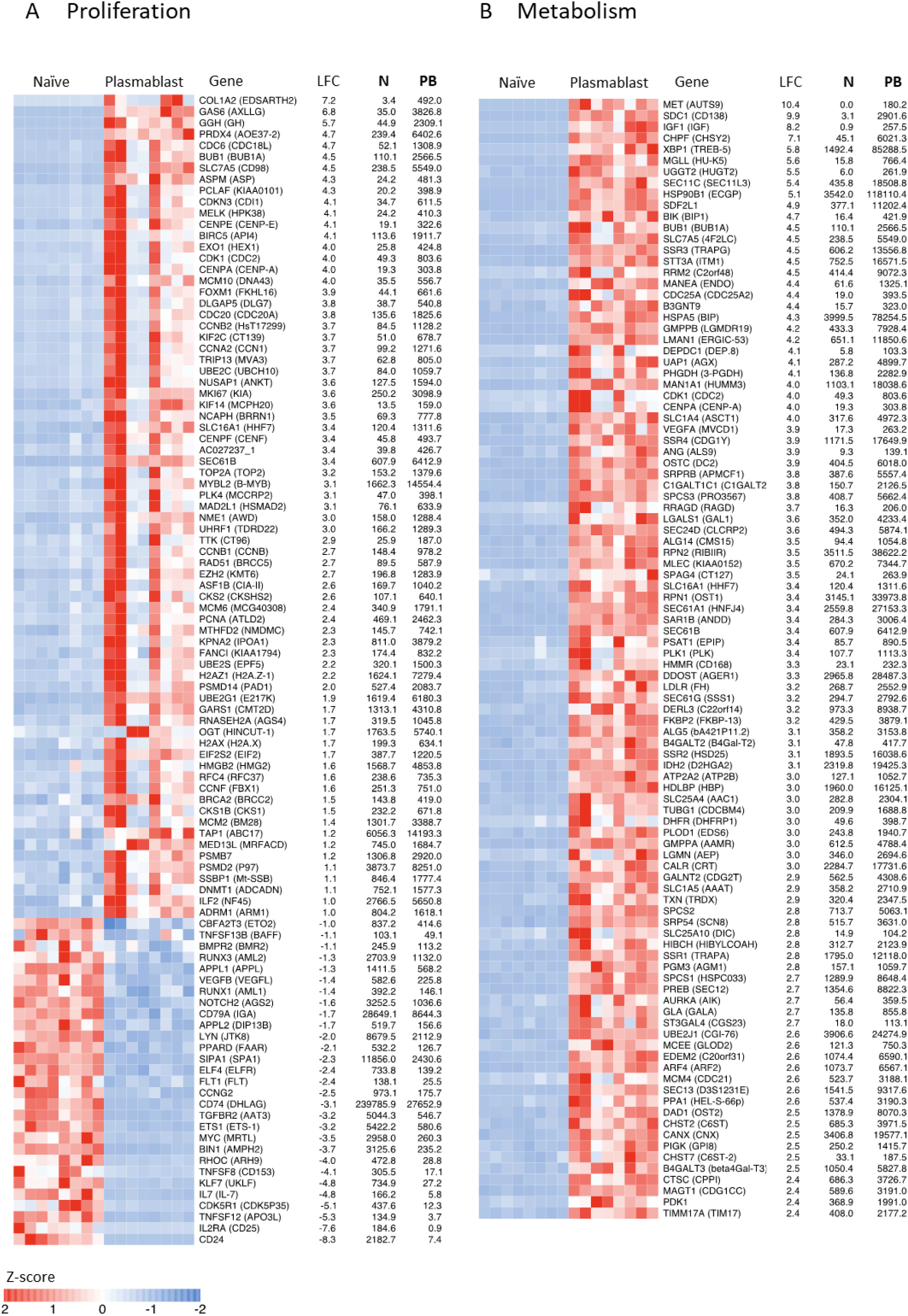
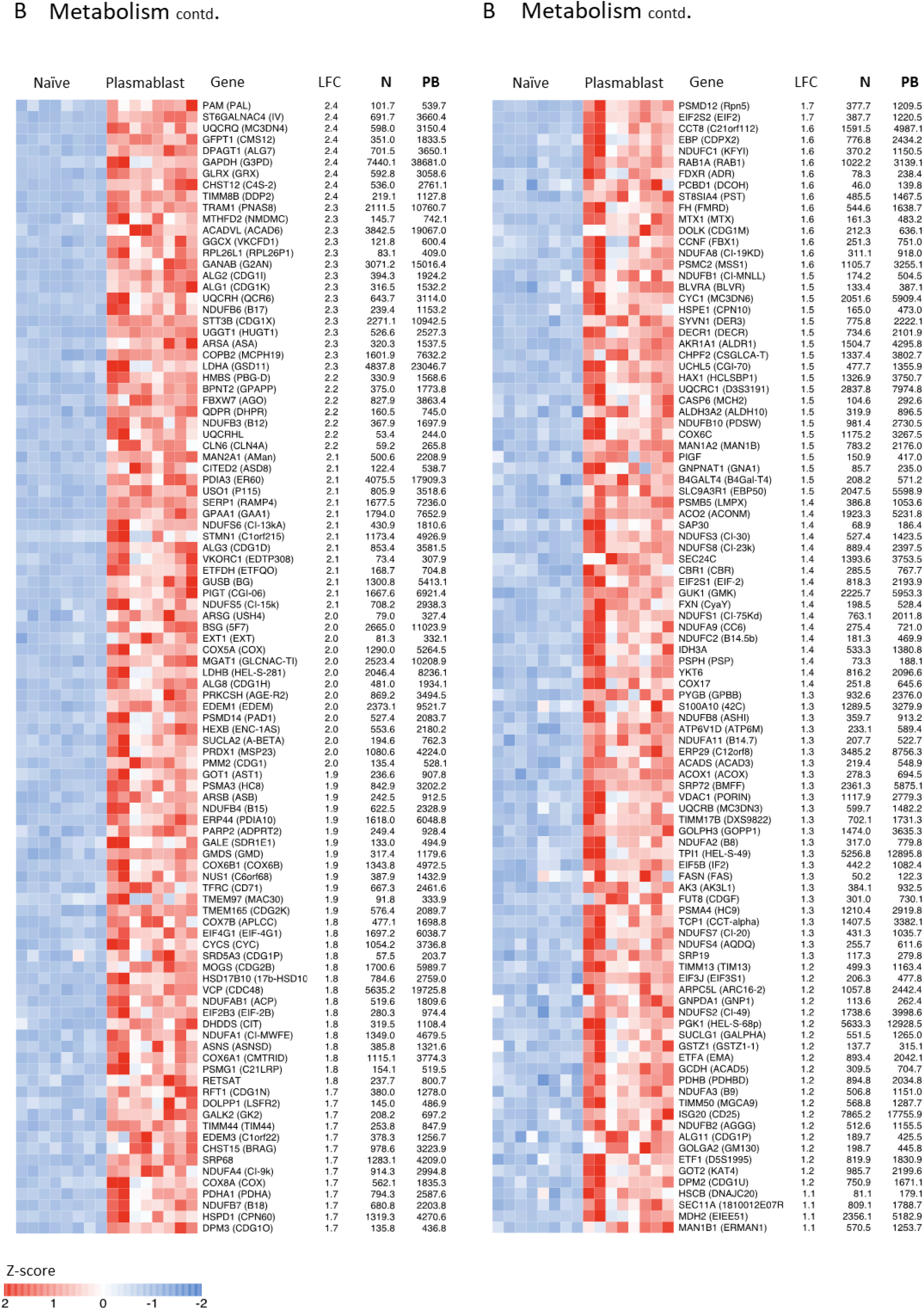
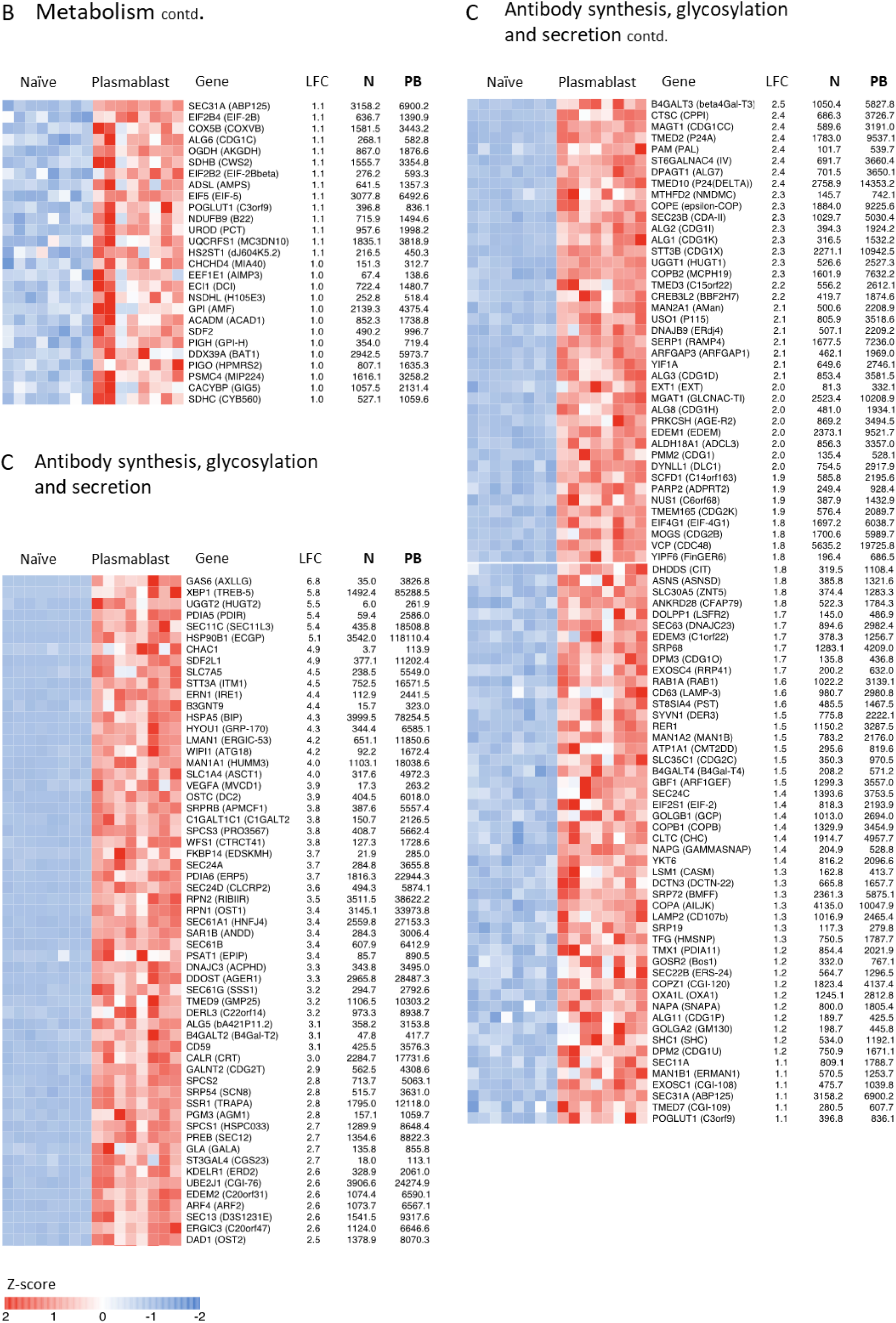
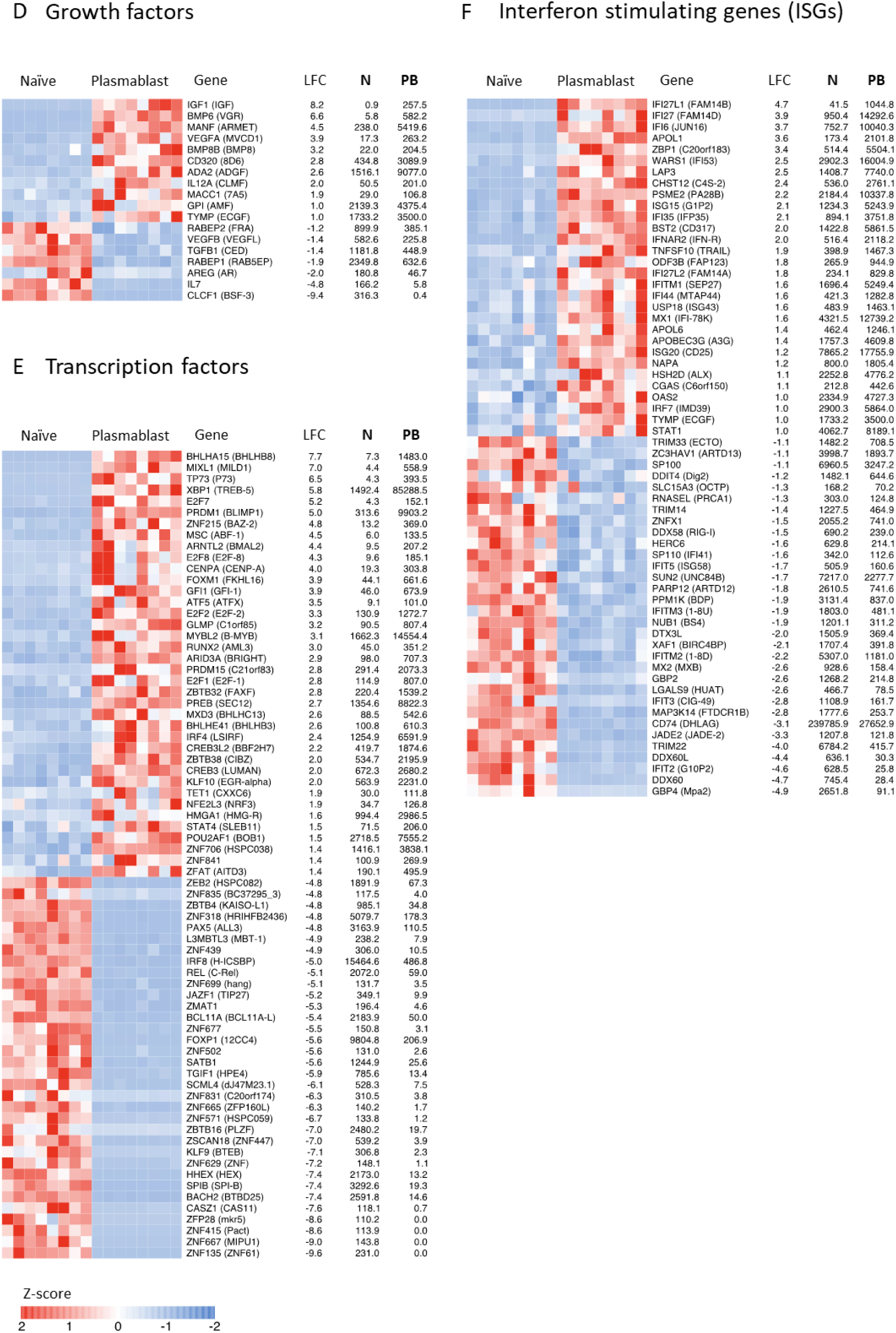
Heat maps of the differentially expressed genes (DEGs) pertaining to select biological processes. Heatmaps (A to F) showing genes related to (A) Proliferation, (B) Metabolism, (C) Antibody synthesis, glycosylation and secretion, (D) Growth factors, (E) Transcription factors (TFs) – top 50 upregulated and top 50 downregulated and (F) Interferon stimulated genes (ISGs). Gradient of high to low gene expression based on z-score normalized counts is indicated from red to blue color. Genes that were significantly differential are shown. Mentioned on right side of heatmap are gene name with common name (in brackets), log2 fold change (LFC), average normalized counts of naïve (N) and average normalized counts of plasmablast (PB).

### Plasmablasts from dengue patients express several cytokines involved in angiogenesis, cellular extravasation and vascular permeability

It is known that B cells produce different cytokines depending on differentiation status (40–43). In contrast to naïve B cells that typically express cytokines such as lymphotoxin B (LTB), TNF- alpha, IL-6 or IL-10 (40–43), emerging studies in murine models show that plasma cells/ plasmablasts can express cytokines such as IL-17, IL10 or IL-35 (44, 45). The expression of these cytokines seems to be context-dependent and appears to have far reaching consequences because mice in which exclusively B cells did not express IL-35 could not recover from experimental autoimmune encephalomyelitis (EAE); but had higher resistance to *Salmonella typhimurium* infection (45). However, virtually nothing is known about cytokines expression by human plasmablasts in dengue infection.

Considering this, we asked what cytokine genes are expressed in human plasmablasts responding to dengue infection. Of the 263 cytokine genes analyzed (46), 33 genes were differentially expressed in plasmablasts compared to naïve B cells. Of these, 15 cytokines were upregulated in plasmablasts. A heatmap of these is shown in **Fig. 7A**. We found that the plasmablasts downregulated key cytokines reported for naïve B cells (*LTB*, -8.0; IL16, -2.4; TGFB1, -1.4; IL32, -1.8 and IL24, -4.8). Although the plasmablasts did not differentially express cytokines such as IL-35 and IL-10 that were previously reported in plasma cells / plasmablasts of murine models (45, 47), we found that they robustly expressed several other cytokines as shown in the heatmap. An extensive list of top 35 biological processes (GO Terms) from STRING (Search Tool for the Retrieval of Interacting Genes/Proteins) associated with these upregulated cytokines is shown in **Fig. 7C** and **Supplementary Table 4.** Many of these upregulated cytokines share functional overlap pointing to vascular permeability, leukocyte extravasation and angiogenesis. Notable among these are *VEGFA* (+3.9), *ADM* (+ 2.3), *ADM2* (+3.3), *BMP6* (+6.6), *IGF1* (+8.2), *MANF* (+4.5) and *TNFSF10* (+1.9), as explained below.

**FIG 7.**
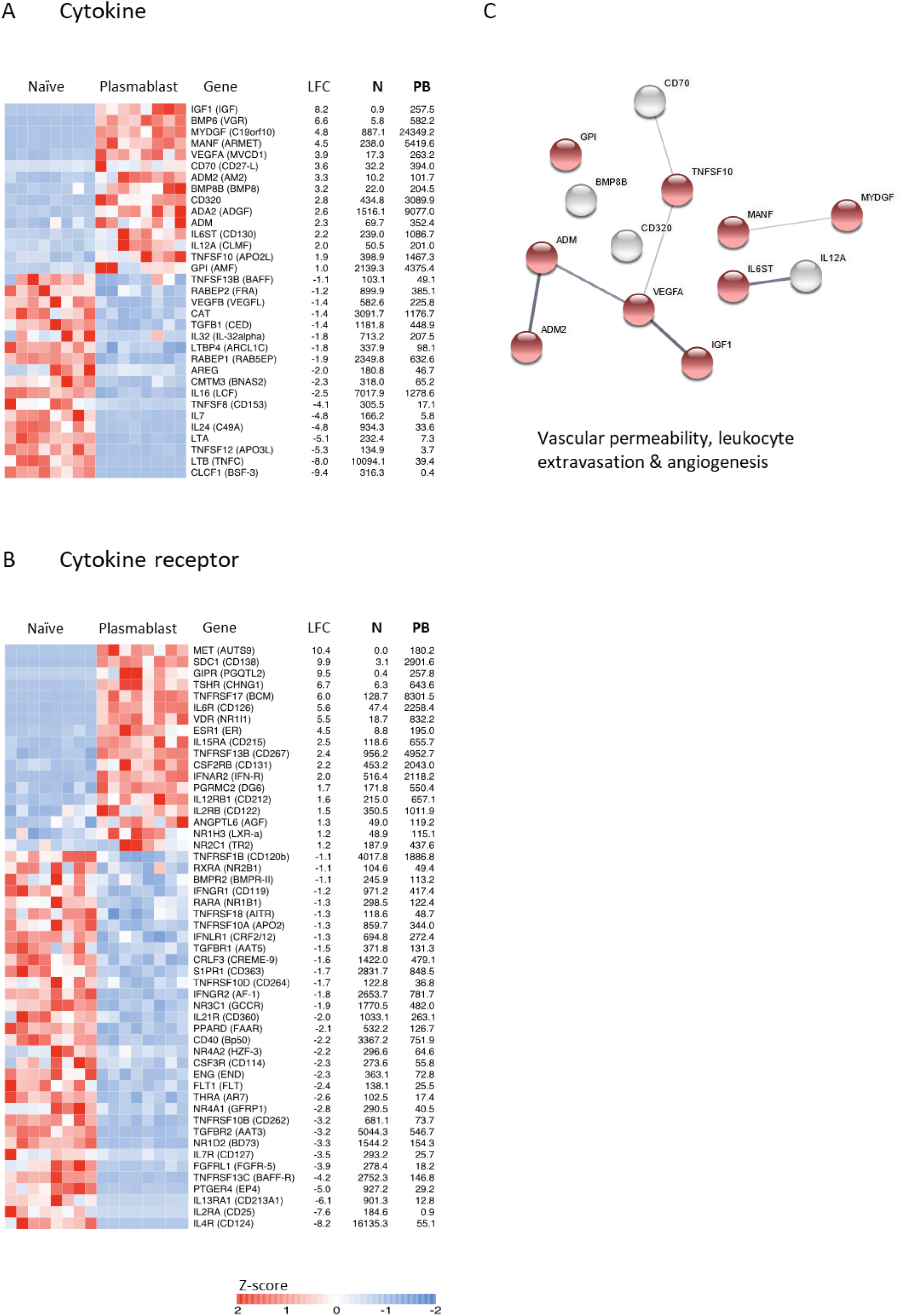
Plasmablasts from dengue patients express several cytokines involved in angiogenesis, cellular extravasation and vascular permeability. Heatmaps showing genes related to (A) cytokines and (B) cytokine receptors. Gradient of high to low gene expression based on z-score normalized counts is indicated from red to blue color. Genes that were significantly differential are shown. Mentioned on right side of heatmap are gene name with common name (in brackets), log2 fold change (LFC), average normalized counts of naïve (N) and average normalized counts of plasmablast (PB). (C) Network analysis performed in STRING describing associations between the upregulated cytokines in plasmablasts based on text-mining; annotated database; experimental evidence; homology. Protein-protein associations are represented by the edges and the thickness of edges corresponds to confidence in the evidences used to deduce the associations (thick – high confidence; narrow – low confidence). Red nodes highlighted represent the cytokines involved in vascular permeability, leukocyte extravasation and angiogenesis.

*VEGFA* is a classic cytokine that was originally named as vascular permeability factor (VPF) and is shown to have multiple effects on endothelial cells including induction of angiogenesis, promotion of vascular permeability and extravasation of leukocytes. *VEGF* is also classically thought to play a pivotal role in mediating plasma leakage in dengue as evidenced by its elevated levels associated with DHF and/or DSS (48–50). *ADM* and *ADM2* are known to synergize *VEGFA* actions on endothelial cells (51, 52). *BMP-6* is known to increase vascular hyperpermeability via distortion of endothelial cell adherence junctions (53). *IGF1* promote endothelial cell migration, tube formation and production of the vasodilator nitric oxide (54, 55). *MANF* is shown to have proangiogenic effects via activation of *VEGF* shared signaling pathways (56). *TNFSF10* (TRAIL) is emerging as a novel angiogenic factor as evidenced by its ability to induce migration of endothelial cell and formation of blood vessel tube similar to *VEGFA* (57). By viewing our data in juxtaposition with these studies it appears that plasmablasts from dengue patients upregulate several cytokine genes involved in vascular permeability, cellular extravasation and angiogenesis.

Interestingly, we observed that plasmablasts robustly upregulated interleukin-6 receptor (*IL-6R,* +5.6) **(Fig. 7B** and **Fig. 1C)** but not *IL6* **(Fig. 7A)**. This indicates that perhaps plasmablasts are more likely to depend on paracrine IL-6 from other cell types (e.g., monocytes or endothelial cells) for their survival. This is consistent with the finding that plasmablasts are engaged in a cross talk with monocytes for expansion in dengue (58). Plasmablasts also upregulated *IFNAR2* (+2.0) **(Fig. 7B)**, which is a receptor for type I IFN. Type I IFN are the most prominent innate cytokines produced in response to viral infections and likely to be higher during the acute febrile phase (59, 60). It is interesting to note that classical studies indicated that type I IFN play an important role in sustaining the survival of effector T and B cells (61–63), suggesting that perhaps like IL-6, type I IFN may also be helping in survival of these plasmablasts in dengue patients. *CD40* (-2.2) was downregulated in plasmablasts, indicating that perhaps these cells are likely to be refractory to T cell help. Additionally, they also down regulated IL-4R (-8.2) indicating that they are also refractory to IL-4 signaling.

### Plasmablasts from dengue patients expressed several adhesion molecules, chemokines and chemokine receptors involved in endothelial interactions and homing to skin or mucosal tissues including intestine

Heatmaps of the differentially expressed adhesion molecules, chemokines and chemokine receptors in plasmablasts compared to naïve B cells is shown in **Fig. 8A, B and C**. An extensive list of top 35 biological processes (GO Terms) associated with these is shown in **Supplementary Table 5**. STRING analysis of the upregulated CAMs, chemokines and chemokine receptors is shown in **Fig. 8D**. We found that many of these upregulated genes share functional overlap pointing to vascular endothelial cell interactions, adhesion, rolling, diapedesis, extravasation and cell migration to inflammatory tissues, skin, mucosa and intestine. Notable among these are discussed below.

**FIG 8.**
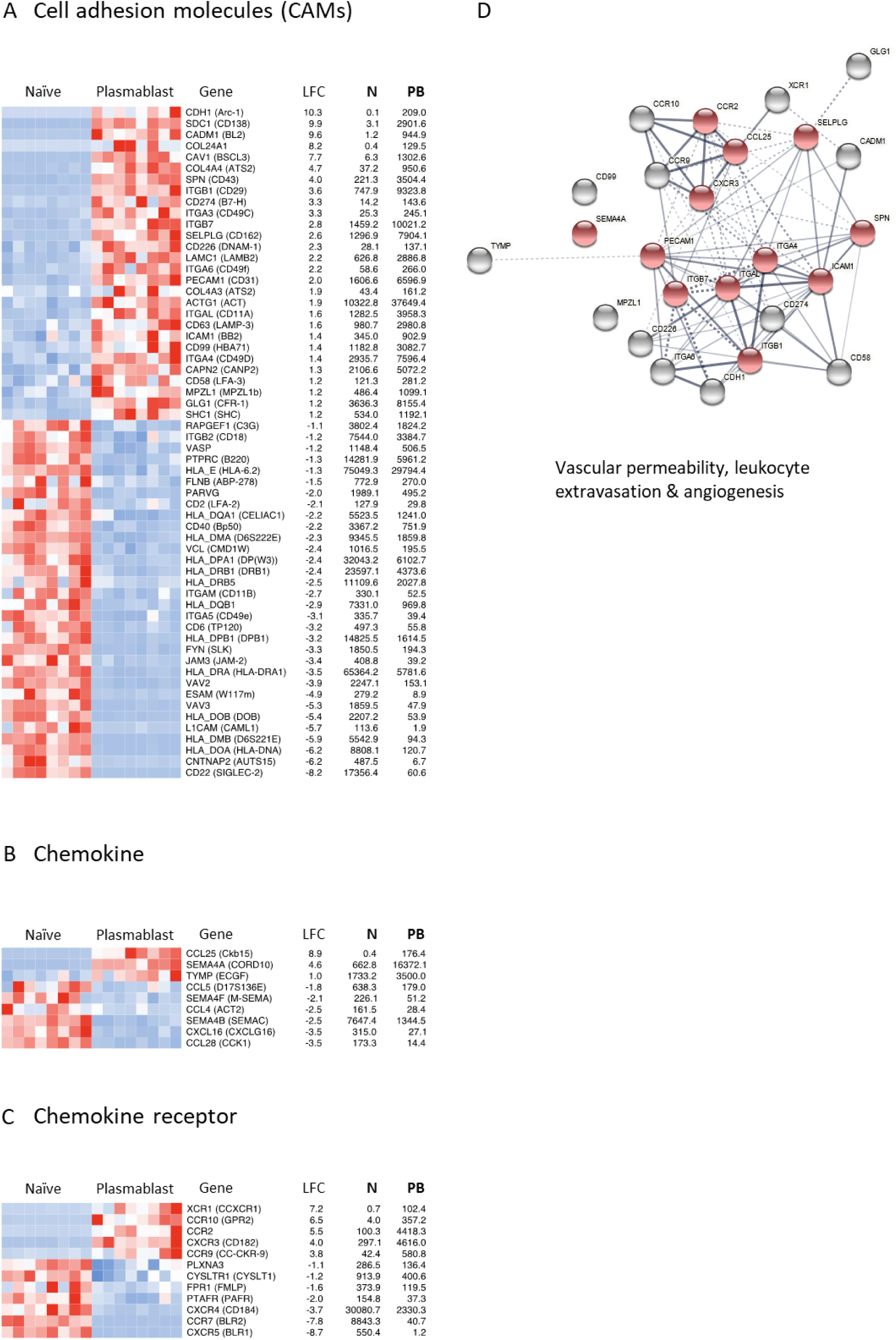
Plasmablasts from dengue patients express several adhesion molecules, chemokines and chemokine receptors involved in endothelial interactions and homing to skin or mucosal tissues including intestine. Heatmaps showing genes related to (A) cell adhesion molecules (CAMs), (B) Chemokine and (C) Chemokine receptors. Gradient of high to low gene expression based on z- score normalized counts is indicated from red to blue color. Genes that were significantly differential are shown. Mentioned on right side of heatmap are gene name with common name in brackets, log2 fold change (LFC), average normalized counts of naïve B cells (N) and average normalized counts of plasmablasts (PB). The genes are arranged as per the descending LFC values. (D) Network analysis performed in STRING describing associations between the upregulated cell adhesion molecules (CAMs), chemokines and chemokine receptors in plasmablasts based on text-mining; annotated database; experimental evidence; homology. Protein-protein associations are represented by the edges and the thickness of edges corresponds to confidence in the evidences used to deduce the associations (thick – high confidence; narrow – low confidence). Red nodes highlighted represent the CAMs, chemokines and chemokine receptors involved in vascular endothelial cell interactions, adhesion and cell migration to inflamed tissues, mucosa and intestine.

Of the upregulated genes associated with adhesion, *CDH1* (E-cadherin, +10.3) is a well-characterized molecule involved in cell-cell adhesion (64, 65). *SDC1/CD138* (+9.9), is known to have pleiotropic functions in cell adhesion (66, 67), homing (25), wound healing (68, 69) and angiogenesis (70, 71). *SDC1* is also known to promote survival of ASC’s in a cell intrinsic manner (72). Cell adhesion molecule-1 (*CADM1*, +9.6) is known to mediate cell-cell adhesion in a calcium independent manner (73). In this regard, it is interesting to note that plasmablasts are likely to be inept generators of calcium flux due to downregulation of CD20 and other calcium flux mediated signaling cascades downstream of BCR as indicated by our GSEA analysis in **Fig. 5F** and thus the cells may be inclined towards calcium independent mechanisms. Caveolin (*CAV1*, +7.7) is a pleiotropic molecule with functions ranging from cell invasion, signaling, intracellular organelle communication and also implicated in mediating endothelial dysfunction by attenuation of nitric oxide under the conditions of hyperlipidemia (74–76). Plasmablasts also showed expression of a wide-spectrum of integrins such as *ITGB1* (+3.6), *ITGA3* (+3.3), *ITGB7* (+2.8), *ITGA6* (+2.2), *ITGAL / CD11a* (+1.6) and *ITGA4/CD49D* (+1.4) that are known to be involved in cell-to-cell or cell-to-extracellular matrix adhesion (77). We also observed a striking decrease in the transcription of a number of several Class II associated genes (*HLA_DPA1, HLA_DRB1, HLA_DRB5, HLA_DQB1, HLA_DOB, HLA_DMB, HLA_DOA*) suggesting that these cells were perhaps incapable of cognate interactions with CD4 T cells.

Examination of chemokines and chemokine receptors showed that the plasmablasts expressed several of them involved in endothelial interactions and homing to skin or mucosal tissues including intestine. As shown in **Fig. 8B and 8C**, plasmablasts expressed CCR9 (+3.8), and CCR10 (+6.5), which are chemotactic receptors for mucosal chemokines such as CCL25/ CCL27/ CCL28. CCL25 is generally expressed on endothelium of small intestine; CCL27 on skin keratinocytes; and CCL28 on a wide variety of mucosal epithelial cells (78–81). Moreover, these cells upregulated CCR2 (+5.5) and CXCR3 (+4.0), the two major chemokine receptors for a variety of chemotactic factors such as CCL2, CCL7, CCL9, CCL10, CCL11, CCL12 that are produced in inflamed tissues. In addition to having equipped with ability to home to skin, mucosal, and intestinal tissues, it is interesting to note that these also expressed XCR1/ CCXCR1/ lymphotactin receptor (+ 7.2) **(Fig. 8C)**. XCR1 is known to bind to lymphotactin / XCL1/ which are emerging as cytokines produced by activated NK cells, NK-T cells and T cells (82, 83). This suggested that, perhaps, these cells are likely to be equipped with ability to migrate towards other activated lymphocytes. Additionally, these cells also expressed semaphorin, SEMA4A (+4.6). Semaphorins and their receptors plays an important role biologically as they function in regulation of immune system, remodeling of ECM and angiogenesis (84–87). Taken together, these findings suggest that plasmablasts from dengue patients are equipped with ability to not only interact with endothelial cells of inflamed tissue, but also produce several mediators involved in vascular permeability, extravasation and angiogenesis, and homing to skin or mucosa as well as intestine

### Plasmablasts from dengue patients and flu vaccinees show qualitatively similar transcriptional profiles

Given that plasmablasts expand massively in dengue patients (5, 22–24) as compared to flu infected / vaccinees (88), we wondered whether the transcription signature we observed in plasmablasts from dengue patients was unique to dengue or could be applied to other situations. Our previous influenza study provided transcriptional profiling of plasmablasts from Flu vaccinees (28) but this study was based on microarray analysis and hence had a restricted gene set analysis. Hence, we retrieved our influenza study samples and performed RNA seq analysis of the plasmablasts versus naïve B cells from three of the flu vaccinees. Similar to what we observed in dengue, plasmablasts from flu vaccinees had a highly distinct transcriptional profile beyond Ig genes compared to naïve B cells. A total of 5008 non-Ig genes were differentially expressed in Flu vaccinee plasmablasts compared to naïve B cells, as shown in volcano plot (**Fig. 9A**). The entire set of the differentially expressed genes is provided in **Supplementary Table 6**. A sample-to-sample correlation analysis of naïve or plasmablast from Dengue patients and naïve or plasmablast from flu vaccinees showed that plasmablasts from Dengue and Flu are strongly correlating (**Fig. 9B**). Comparison of median difference in z-scores (between plasmablast and naïve) of differentially expressed genes showed a positive, significant and very large correlation in dengue and flu (rho = 0.95, p value < .001). Additionally, a majority of GSEA terms for flu plasmablast versus naïve showed normalized enrichment scores similar to that of dengue specific terms (**Fig. 9C**). Similar to dengue, many of the cytokine and adhesion molecules involved in vascular permeability, extravasation and angiogenesis are also upregulated in plasmablasts of the flu vaccinees (**Fig. 9D and 9E**).

**FIG 9.**
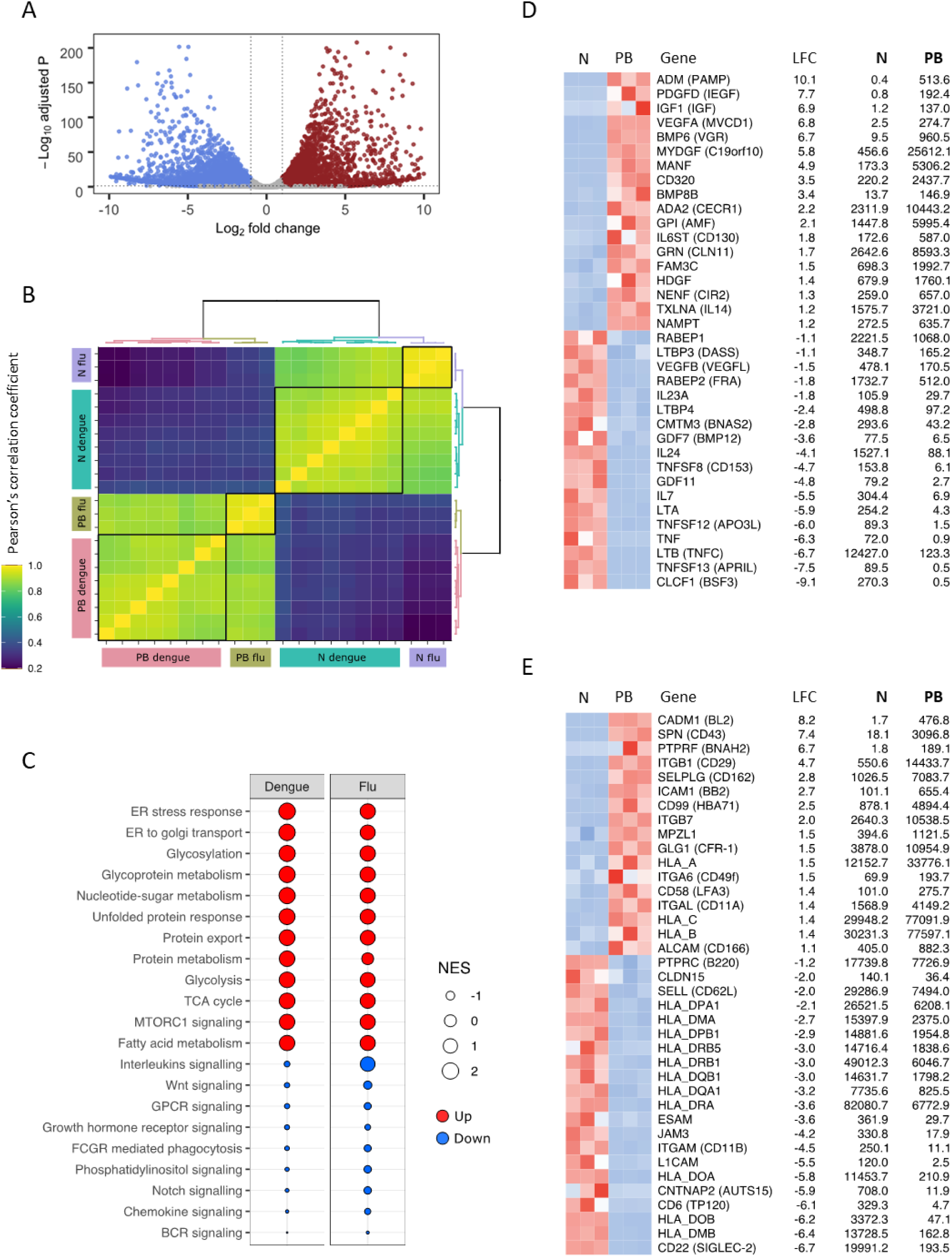
Transcriptional profile of plasmablast derived from flu vaccinees and their comparison with plasmablast derived from Dengue subjects. (A) Volcano plot with -log (adjusted P value) on y-axis and log2 (fold change) on x-axis. Scattered dots represent genes. Red dots are genes that are significantly upregulated genes and blue dots represent significantly downregulated genes in plasmablasts versus naïve B cells (total: 5008 genes), whereas grey dots are either non-significant, non-differential or both. (for *significant*: adjusted P value < 0.05; for *differential*: log2 fold change >= 1 or =<-1). (B) Sample-to-sample correlation heatmap depicting Pearson’s correlation coefficient among samples of dengue and flu. Grouping patterns are represented by agglomerative hierarchical clustering by using Ward.D2 method. Color key depicts a strong correlation for values more than 0.8 (green) or weak correlation for values less than 0.4 (blue). (C) Comparison of GSEA terms enriched for plasmablast versus naïve in Flu with the terms enriched for plasmablast versus naïve in Dengue. Size of the bubble is proportional to the normalized enrichment scores (NES), red and blue color represents terms with positive and negative NES, respectively. Heatmaps showing genes related to (D) Cytokine, (E) Cell adhesion molecules (CAMs). Gradient of high to low gene expression based on z-score normalized counts is indicated from red to blue color. Genes that were significantly differential are shown. Mentioned on right side of heatmap are gene name with common name in brackets, log2 fold change (LFC), average normalized counts of naïve B cells (N) and average normalized counts of plasmablasts (PB). The genes are arranged as per the descending LFC values.

### Plasmablast responses in primary versus secondary dengue infections with different grades of disease severity

Although the results presented in the above sections showed that the transcriptional profiles of the plasmablasts from patients with primary and secondary dengue infection or patients with DW and SD did not cluster, plasmablasts upregulated several genes involved in vascular permeability, the hallmark feature of dengue. Hence, we wondered whether the extent of the plasmablast expansion differed depending on primary and secondary dengue infection status or disease severity. We found that the plasmablast expansion was relatively lower in patients with primary dengue infection as compared to secondary dengue (**Fig. 10A**). The expansion was also relatively lower in DI cases as compared to DW/SD cases **(Fig. 10B).** Even though the plasmablast expansion was lower in the primary dengue patients’ group as a whole compared to secondary dengue patients, within this primary dengue group, the expansion was even lower in patients with mild dengue infection (DI) as compared to dengue patients with warning signs (DW) or severe dengue (SD) (**Fig. 10C)**.

**FIG 10.**
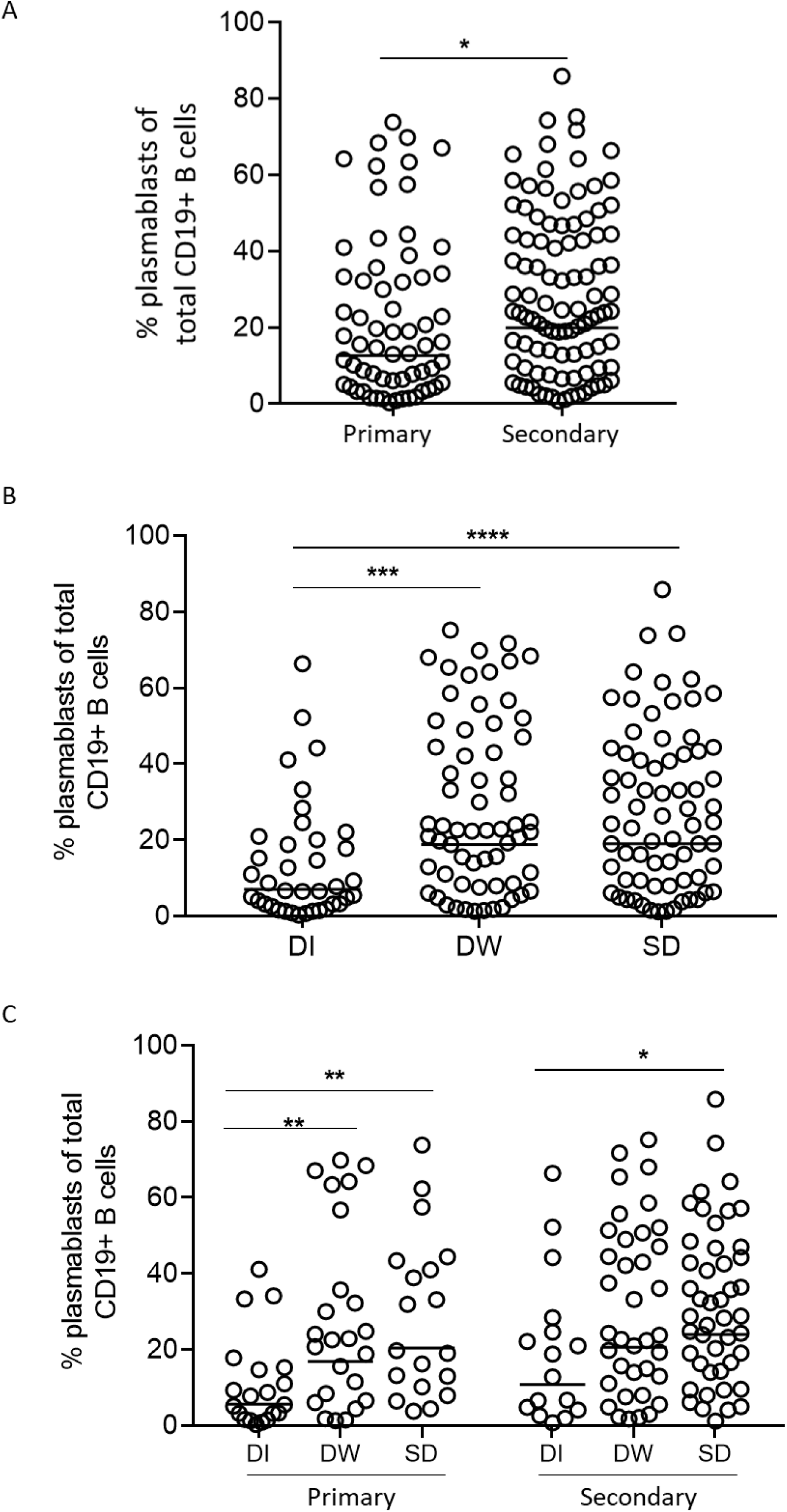
Plasmablast response in primary/secondary dengue patients and patients with different disease severity. (A) Scatter dot plot showing plasmablast frequencies of total CD19+ B cells in primary (n=63) and secondary (n=99) dengue patients. The box represents the geometric mean of all points. (B) Scatter dot plot showing plasmablast frequencies of total CD19+ B cells in patients with dengue infection (DI) (n=36), dengue with warning signs (DW) (n=61) and severe dengue (SD) (n=65). (C) Scatter dot plot showing plasmablast frequencies of total CD19+ B cells in patients with dengue infection (DI), dengue with warning signs (DW) and severe dengue (SD) among primary and secondary dengue patients. The line represents the geometric mean of all points.

## Discussion

This is the first detailed characterization of plasmablasts from dengue patients in India. Our study provides a detailed understanding of the plasmablast expansion, phenotypes, functions and gene expression profiles in dengue patients. Consistent with studies from other parts of the world, we show that plasmablasts expand massively during dengue febrile illness. We found that this expansion was significantly lower in patients with primary dengue infections compared to patients with secondary dengue infections; and significantly lower in DI cases compared to DW/SD cases. We show that these massively expanding plasmablasts expressed several markers of inflammatory tissue/ mucosal homing and upregulated several cytokine genes that are involved in vascular permeability/ angiogenesis.

While our observation that plasmablasts expressed several markers of inflammatory tissue/ mucosal homing is not unexpected for a systemic infection such as dengue (78, 89), we were surprised to learn that plasmablasts from dengue patients also expressed several cytokines that are involved in vascular permeability/ angiogenesis (48–56). Some of these (e.g., VEGFA) are well known endothelial permeability factors (90–92). Increased levels of plasma VEGFA are known to be associated with dengue hemorrhage or shock (48–50). While many of these vascular permeability factors can also be produced by other innate cell types (93, 94), it remains to be determined whether the plasmablasts from dengue patients constitutively secrete these cytokines or secrete them only in a localized manner when interacting with endothelial cells, and if so, what could be the potential physiological effects of the plasmablast- endothelial interactions on plasma leakage.

It is interesting to note from our study that while these transcriptional profiles were not strikingly different in plasmablasts derived from dengue patients with primary/ secondary infections or DW/ SD disease grades or even from flu vaccinees, the level of expansion could differ depending on the context. Usually, plasmablast expansion tend to be minimal (in the range of 1-7% of the B cells) in flu vaccines (88). By contrast, plasmablast show a wide range of expansion in dengue patients, with frequencies reaching as high as 80% of the total B cell population in some patients– making up nearly 30% of the total PBMC. While the expansion was relatively lower in primary dengue patients as a whole group, even within this primary group, we found that DW/SD cases had a much higher expansion compared to DI cases. Interestingly, in situations of other viral infections that cause hemorrhagic fever during primary infection (e.g., Ebola) a massive expansion of the plasmablasts has been reported (28, 95). Taken together, these findings raise an interesting question on whether a massive expansion of plasmablasts and their coordinated expression of a multitude of immune mediators that are potentially involved in endothelial interactions and vascular permeability could be a common feature of hemorrhagic fever viruses in general. These findings highlight the need for further studies to carefully examine the precise role of plasmablast responses in hemorrhagic fevers.

## Methods

### Dengue patient recruitment

The clinical site for this study is Department of pediatrics of the All India Institute of Medical Sciences (AIIMS), New Delhi, India. Patients in this study were enrolled between the years 2012-2019. The study was approved by the institutional ethical committee, and informed consent/ assent was obtained from the study participants prior to recruitment to the study. Patients included in the study were negative for malaria antigen, and chikungunya IgM.

### Dengue confirmation

Dengue virus infection was confirmed by a combination of methods, including reverse transcription (RT)-PCR as described previously (9), NS1 Elisa (Enzyme linked immunosorbent assay) (J Mitra) and/or dengue IgM/IgG Elisa (PanBio).

### Disease classification

Based on extensive clinical laboratory tests and evaluation, the attending physicians at each clinical site classified the dengue disease grade based on the WHO 2009 guidelines as dengue infection without warning signs (DI), dengue with warning signs (DW) and severe dengue (SD) (96) at the time of recruitment.

### PBMC and plasma isolation

Blood sample was collected in Vacutainer CPT tubes (Becton Dickenson, Cat# 362761). The PBMC’s and plasma were separated as described previously (22). The PBMC’s were washed extensively in 1% complete RPMI (RPMI containing 1% FCS, 1X Pen/Strep, 1X Glutamine) and used for analytical flow cytometry and *ex vivo* Dengue-specific ELISPOT assays.

### Characterization of primary and secondary infections

Primary and secondary infections were classified using the standard Panbio capture ELISAs (Alere, 01PE10/01PE20) that quantify IgM and IgG ratios in plasma diluted at 1:100. As per the WHO criteria, primary dengue infections were defined as those who were confirmed as dengue infected by NS1 and/or PCR but failed to induce any immunoglobulin response or in whom only IgM was induced or both IgM and IgG were induced but with the IgM/IgG ratio being >1.2. Secondary infections were defined as those who were confirmed as dengue infected by NS1 and/or PCR and in whom only IgG was induced or both IgM and IgG were induced with IgM/ IgG ratio being <1.2 (96).

### Flow cytometry staining and analysis

Extensively washed PBMC’s were stained with fluorochrome labeled antibodies and fixable viable dye eFluor 780 (eBioscience, Cat# 65-0865- 14) for live dead exclusion. For plasmablast analysis, cells were stained with CD19 (Biolegend, Cat# 302252), CD20 (Biolegend, Cat# 302312), CD38 (BD, Cat# 555460) and CD27 (BD, Cat# 563092) and acquired on a BD FACS Canto II. Plasmablast staining was analyzed using Flow Jo software (TreeStar Inc) starting with a lymphocyte gate that included the blasting cells and plasmablasts were described as cells that are CD3-, CD20-, CD19+/Int, CD27+ and CD38+. The plasmablasts were further stained with Ki67 (BD, Cat# 561283), CD71 (BD, Cat# 555537), IgM (Biolegend, Cat# 314512), IgG (BD, Cat# 561298), IgD (BD, Cat# 555778), CXCR5 (R&D, Cat# FAB190P), CXCR4 (BD, Cat# 555974), CXCR3 (BD, Cat# 560831), CD24 (BD, Cat# 555427), CCR7 (BD, Cat# 560765), CCR2 (R&D, Cat# FAV151P), CLA (Biolegend, Cat# 321312), BAFFR (Biolegend, Cat# 316916), IL-6R (BD, Cat# 551850), and CCR9 (Biolegend, Cat# 358907) for their phenotype. For intracellular protein staining, cells were permeabilized with Cytofix/Cytoperm buffer (BD), followed by staining for 60 min with antibodies that were diluted in Perm/Wash buffer (BD, Cat# 554723;)

### Dengue-specific ELISpot assay

Dengue-specific ELISpot assay was performed on PBMCs isolated as previously described (22). Briefly, 96-well ELISpot plates (Millipore, Cat# MSIP45) were coated overnight with either Dengue 2 fixed virus antigen (Microbix, Cat# EL-22-02) or polyvalent total immunoglobulin (Invitrogen, Cat# H17000). Before adding PBMC’s, the plates were washed thoroughly and blocked for 2 hrs at 37°C with RPMI containing 10% FBS. Extensively washed 0.1X10^6^ PBMC’s were then added to the plate and diluted 3-fold till the last well had a cell number of 45 cells and incubated overnight at 37°C undisturbed. The cells were removed; plates were washed with 1X PBS+0.05% Tween 20 and developed with Goat anti- human IgG Biotin (eBioscience, Cat# 13-4998-83), IgM Biotin (Sigma, Cat# B1265) and Avidin HRP (eBioscience, Cat# 18-4100-51). The spots were developed with AEC substrate (BD, Cat# 551951) and scanned using an automated ELISpot counter (CTL, Cellular Technologies LTD).

### Fluorescence-activated cell sorter (FACS) sorting of naïve B cells and plasmablasts

PBMCs isolated from peripheral whole blood of dengue patients were stained with the relevant antibodies at 4°C for 30 min, washed thoroughly, suspended in PBS containing 2% FCS, and sorted on a BD FACS Aria III (BD) with high forward-scatter gates to account for the larger blasting effector lymphocytes. CD19+ CD20+ IgD+ CD71- CD27- CD38- naïve B cells and CD19+/int CD20- IgD- CD71+ CD27+ CD38+ plasmablasts were isolated to a purity of more than 90%.

### RNA isolation and Library preparation

The sorted cells were washed thoroughly and the cell pellet was suspended in RLT Buffer (Qiagen) and stored at -80^0^C until RNA extraction. All the samples were processed simultaneously for RNA extraction. RNA was isolated from sorted B cells for each patient using the RNeasy Micro kit (Qiagen) with on-column DNase digestion. RNA quality was assessed using an Agilent Bioanalyzer and 500pg of total RNA was used as input for cDNA synthesis using the Clontech SMART-Seq v4 Ultra Low Input RNA kit (Takara Bio) according to the manufacturer’s instructions. Amplified cDNA was fragmented and appended with dual-indexed bar codes using the Nextera XT DNA Library Preparation kit (Illumina). Libraries were validated by capillary electrophoresis on an Agilent 4200 TapeStation, pooled at equimolar concentrations, and sequenced on an Illumina HiSeq3000 at 100SR, yielding approximately 20 million reads per sample.

### RNA-seq data analysis

Sequenced reads were mapped to human genome GRCh38. FastQC was used to monitor the quality of sequences. Adaptor sequences were trimmed. Quantification of transcripts was done by using the featurecounts option of the Subread package. Raw counts from 60,448 genes were produced as an output from which 407 immunoglobulin genes were removed for further analysis. Differential expression analysis was performed with total 60,041 genes using *DESeq2 package* (97). Low count genes (45, 268) with FPKM < 10 in more than five samples were filtered out. Among the 14,773 genes, those with adjusted P value (Benjamini-Hochberg method) < 0.05 and p value < 0.01, were declared as significant. Log2 fold change of more than or equal to +1 was used as threshold to declare genes as upregulated whereas log2 fold change of less than or equal to -1 was used for assigning downregulated genes. Further, genes with low average normalized counts of 100 or less in both conditions (plasmablast and naïve) were excluded from the differentially expressed gene list. PCA plots were generated from the 14,773 genes. For heatmaps, normalized counts transformed to their z-scores were used. Hierarchical clustering was created using Euclidean distance with Ward.D2 method. All analysis was performed in R studio software using *DESeq2*, *pheatmap*, *dendextend*, *dplyr*, *reshape2*, *ggplot2*, biomaRt, *prcomp* and *rgl* packages.

### Gene Set Enrichment Analysis (GSEA)

GSEA software v 4.0.3 from Broad institute (98, 99) was used for the enrichment analysis. Our expression dataset was run against Hallmark, KEGG pathway, Reactome pathway and GO Biological process gene sets from the Broad Institute’s Molecular signature database (MSigDB). 1000 gene set permutations were used to estimate the nominal P value of enrichment score (ES). To account for gene set size the ES was normalized to get a normalized enrichment score (NES) for each term. False discovery rate (FDR) was used to account for multiple testing, and terms with FDR < 25% were considered.

### Protein Network Analysis

Protein-protein associations among the upregulated genes in cytokines, cell adhesion molecules, chemokines and chemokine receptors were calculated using STRING software v11.0.(100). Each direct and indirect association was scored based on experimental evidence, curated database, text-mining, co-expression and protein homology. Functional overlap of the genes was analyzed based on the functional enrichment of biological process.

### Correlation analysis

For sample-to-sample correlation analysis of dengue and flu subjects, log of normalized counts of total 4126 differentially expressed genes in dengue plasmablast versus naïve were used for calculating Pearson’s correlation coefficient. Heatmap was made using *heatmaply* package in R studio. Ward.D2 method was used for clustering. Further, to calculate the overall correlation between dengue and flu, differentially expressed genes in dengue were used and compared their expression profile in Flu. Pearson’s correlation coefficient was calculated using the median z-score difference between plasmablast and naïve of dengue and flu. Those with positive and negative median z-score difference were assigned as upregulated and downreuglated genes, respectively.

### Statistical Analysis

All data was tabulated using MS Excel and analyzed using GraphPad prism software. For analysis of groups, unpaired two-tailed t test was used to determine statistical significance and p values were interpreted as * p≤0.05; ** p≤0.01; *** p≤0.001, ****p=<0.0001.

### Data availability

The dengue RNA seq dataset is deposited in Gene expression omnibus (GEO) with the accession code- GSE171487.

## Acknowledgements

This work was supported by U.S. National Institutes of Health grant ICIDR 1UO1A/115654; NIH- DBT, Human immunology Project Consortium (HIPC) grant BT/PR30260/MED/15/194/2018; Government of India Department of Biotechnology grant DBT BT/PR5132/MED/15/85/2012; We thank Aditya Rathi (ICGEB-TACF) for FACS sorting of the PBMCs. We are very thankful to Dr. Steven Bosinger and Dr. Kathryn Pellegrini, Yerkes Genomics Core, Emory University, Atlanta, Georgia, USA for RNA sequencing of the samples. The authors thank Mr. Satendra Singh, Mr. Ajay Singh, ICGEB, New Delhi for technical support.

---

**Supplementary Table 1** List of all the differential genes between plasmablasts vs naïve from dengue patients. (provided as an excel file)

**Supplementary Table 2** List of enriched gene sets between plasmablasts vs naïve from dengue patients. (provided as an excel file)

**Supplementary Table 3** List of genes and their log2 fold change involved in the selected enriched gene sets between plasmablasts vs naïve from dengue patients. (provided as an excel file)

**Supplementary Table 4** List of top pathways associated with upregulated cytokines as highlighted by STRING (provided as an excel file)

**Supplementary Table 5** List of top pathways associated with upregulated CAMs, chemokines and chemokine receptors as highlighted by STRING (provided as an excel file)

**Supplementary Table 6** List of all the differential genes between plasmablasts vs naïve from flu vaccinees. (provided as an excel file)

